# SARS-CoV-2 infected cells present HLA-I peptides from canonical and out-of-frame ORFs

**DOI:** 10.1101/2020.10.02.324145

**Authors:** Shira Weingarten-Gabbay, Susan Klaeger, Siranush Sarkizova, Leah R. Pearlman, Da-Yuan Chen, Matthew R. Bauer, Hannah B. Taylor, Hasahn L. Conway, Christopher H. Tomkins-Tinch, Yaara Finkel, Aharon Nachshon, Matteo Gentili, Keith D. Rivera, Derin B. Keskin, Charles M. Rice, Karl R. Clauser, Nir Hacohen, Steven A. Carr, Jennifer G. Abelin, Mohsan Saeed, Pardis C. Sabeti

## Abstract

T cell-mediated immunity may play a critical role in controlling and establishing protective immunity against SARS-CoV-2 infection; yet the repertoire of viral epitopes responsible for T cell response activation remains mostly unknown. Identification of viral peptides presented on class I human leukocyte antigen (HLA-I) can reveal epitopes for recognition by cytotoxic T cells and potential incorporation into vaccines. Here, we report the first HLA-I immunopeptidome of SARS-CoV-2 in two human cell lines at different times post-infection using mass spectrometry. We found HLA-I peptides derived not only from canonical ORFs, but also from internal out-of-frame ORFs in Spike and Nucleoprotein not captured by current vaccines. Proteomics analyses of infected cells revealed that SARS-CoV-2 may interfere with antigen processing and immune signaling pathways. Based on the endogenously processed and presented viral peptides that we identified, we estimate that a pool of 24 peptides would provide one or more peptides for presentation by at least one HLA allele in 99% of the human population. These biological insights and the list of naturally presented SARS-CoV-2 peptides will facilitate data-driven selection of peptides for immune monitoring and vaccine development.

## INTRODUCTION

As researchers work to develop effective vaccines and therapeutics for Severe Acute Respiratory Syndrome Coronavirus 2 (SARS-CoV-2), the virus causing the ongoing Coronavirus Disease 19 (COVID-19) pandemic (Lu et al., 2020), it will be critical to decipher how infected host cells interact with the immune system. Previous insights from SARS-CoV and MERS-CoV, as well as emerging evidence from SARS-CoV-2, imply that T cell responses play an essential role in SARS-CoV-2 immunity and viral clearance (Altmann and Boyton, 2020; Grifoni et al., 2020a; Le Bert et al., 2020; Moderbacher et al., 2020; Sekine et al., 2020). When viruses infect cells, their proteins are processed and presented on the host cell surface by class I human leukocyte antigen (HLA-I). Circulating cytotoxic T cells recognize the presented foreign antigens and initiate an immune response, resulting in the clearance of infected cells. Investigating the repertoire of SARS-CoV-2 derived HLA-I peptides will enable identification of viral signatures responsible for activation of cytotoxic T cells.

The vast majority of studies that have interrogated the interaction between T cells and SARS-CoV-2 antigens to date utilized bioinformatic predictions of HLA-I binding affinity (Campbell et al., 2020; Grifoni et al., 2020b; Nguyen et al., 2020; Poran et al., 2020a). While HLA-I prediction is undoubtedly a useful tool to identify putative antigens, it has limitations. First, antigen processing and presentation is a multi-step biological pathway, that includes translation of source proteins, degradation by the proteasome, peptide cleavage by aminopeptidases, peptide translocation into the endoplasmic reticulum (ER) by transporter associated with antigen presentation (TAP) and finally, HLA-I binding (Neefjes et al., 2011). Although many computational predictors now account for some of these steps, the average positive predictive values achieved across HLA alleles is still ~64%(Sarkizova et al., 2020). Second, prediction models do not account for ways in which viruses may manipulate cellular processes upon infection, which can affect antigen presentation. For example, viruses can attenuate translation of host proteins, downregulate the proteasome machinery, and interfere with HLA-I expression (Hansen and Bouvier, 2009; Sonenberg and Hinnebusch, 2009). These changes shape the collection of viral and human-derived HLA-I peptides presented to the immune system. Third, prediction models do not capture subcellular localization of SARS-CoV-2 proteins. While it is unclear if and how this localization might affect antigen presentation, it is known that many of the viral non-structural proteins (nsps) are localized to the replicase-transcriptase complex (RTC) in double-membrane vesicles (DMVs) while the structural proteins, like S, E, and M, are localized to the ER and move along the secretory pathway (Astuti and Ysrafil, 2020; Fehr and Perlman, 2015). Thus, experimental measurements of naturally presented peptides is essential to our understanding of SARS-CoV-2 immune response.

Mass spectrometry based HLA-I immunopeptidomics provides a direct and unbiased method to discover endogenously presented peptides and to learn rules that govern antigen processing and presentation(Abelin et al., 2017; Bassani-Sternberg and Gfeller, 2016; Chong et al., 2018; Sarkizova et al., 2020). This technology has facilitated the detection of tumor specific antigens in a wide range of cancers (Bassani-Sternberg et al., 2016; Bilich et al., 2019; Schuster et al., 2017), and identified virus-derived HLA-I peptides for a handful of viruses including West Nile virus, vaccinia virus, human immunodeficiency virus (HIV), human cytomegalovirus (HCMV) and measles virus (Croft et al., 2013; Erhard et al., 2018; McMurtrey et al., 2008; Rucevic et al., 2016; Schellens et al., 2015a; Ternette et al., 2016). These infectious disease studies revealed new antigens, characterized the kinetics of presented peptides during infection, and identified the sequences that activate T cell responses. Leveraging HLA-I immunopeptidome datasets to learn virus specific antigen processing rules will improve our ability to accurately predict viral epitopes and utilize them to study immune responses in COVID-19 patients.

While unbiased relative to presentation, analysis of mass spectrometry-derived data relies upon selecting a set of viral ORFs to interrogate, and has largely focused on canonical ORFs. Over the past decade, genome-wide profiling of translated sequences has revealed a striking number of non-canonical ORFs in mammalian and viral genomes (Finkel et al., 2020a; Ingolia et al., 2009, 2011; Stern-Ginossar et al., 2012). While the function of most of these non-canonical ORFs remains unknown, it is becoming clear that the translated polypeptides, including from short ORFs, serve as fruitful substrates for the antigen presentation machinery (Chen et al., 2020; Ingolia et al., 2014; Ouspenskaia et al., 2020; Starck and Shastri, 2016). Importantly, a recent study identified 23 novel ORFs in the genome of SARS-CoV-2, some of which have higher expression levels than the canonical viral ORFs (Finkel et al., 2020b). Whether these non-canonical ORFs give rise to HLA-I bound peptides remains unknown.

Here, we present the first comprehensive examination of HLA-I immunopeptidome in two SARS-CoV-2-infected human cell lines, and complement this analysis with RNA-seq and global proteomics measurements. We identify viral HLA-I peptides that are derived from both canonical and non-canonical ORFs and monitor their dynamics over multiple timepoints post infection. Using whole proteome measurements, we show that SARS-CoV-2 interferes with the cellular proteasomal pathway, potentially resulting in lower presentation of viral peptides. The list of endogenously processed SARS-CoV-2 peptides presented are estimated to cover at least one HLA-I allele in 99% of the human population. Our study can inform future T cell assays in patients and aid in the design of efficacious vaccines.

## RESULTS

### Profiling HLA-I peptides in SARS-CoV-2 infected cells by mass spectrometry

To interrogate the repertoire of human and viral HLA-I peptides, we immunoprecipitated (IP) HLA-I proteins from SARS-CoV-2-infected human lung A549 cells and human kidney HEK293T cells that were transduced to stably express ACE2 and TMPRSS2, two known viral entry factors. We then analyzed their HLA bound peptides by liquid chromatography tandem mass spectrometry (LC-MS/MS) (**Fig. 1A**). We also analyzed the whole proteome of the IP flowthrough by LC-MS/MS and performed RNA-seq to determine viral abundances and to examine the effect of SARS-CoV-2 on human gene expression. To investigate the kinetics of HLA-I presentation, we performed a time course experiment at 12h, 18h and 24h post-infection in both cell lines. To allow for the detection of peptides from the complete translatome of SARS-CoV-2, we combined the 23 novel ORFs recently identified by Finkel *et al*. (Finkel et al., 2020b) with the list of canonical ORFs and the human refseq database for LC-MS/MS data analysis (**Fig. 1A**).

**Figure 1.**
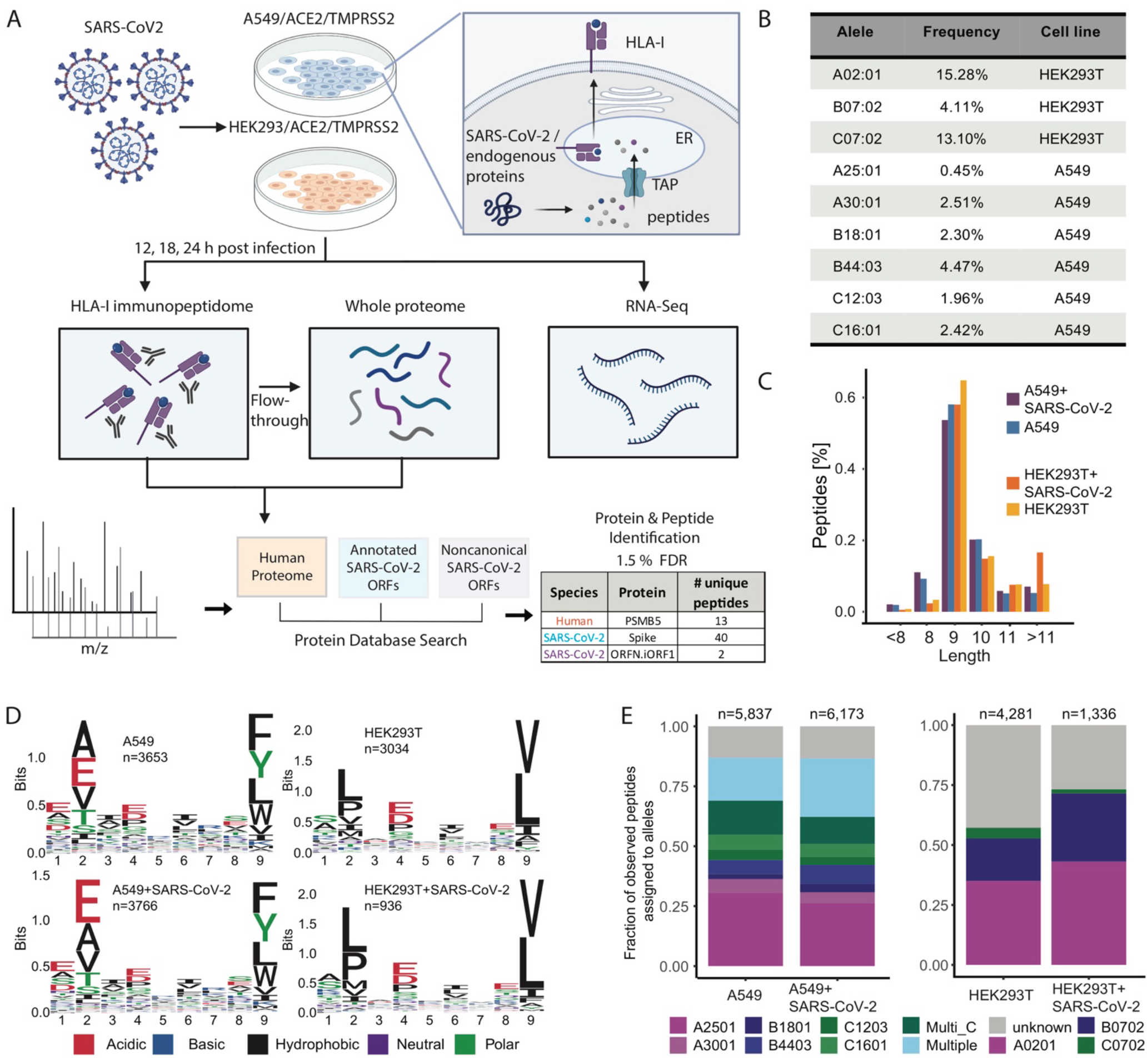
Experimental design and measurements of HLA-I immunopeptidome, whole proteome and RNA-seq in SARS-CoV-2 infected cells. **(A)** Schematic representation of the experiment and the antigen presentation pathway. A549/ACE2/TMPRSS2 and HEK293T/ACE2/TMPRSS2 cells were infected with SARS-CoV-2 (Washington strain, accession number MN985325) at Multiplicity of Infection (MOI) of 3 in a BSL3 facility and harvested at 12, 18 and 24hpi (hours post infection). For HLA-I immunopeptidome measurements cells were lysed and HLA-I peptide complexes were immunoprecipitated. The flow-through after immunoprecipitation was collected for whole proteome measurements. For RNA-seq we added Trizol to infected cells, purified the RNA and performed strandspecific short reads sequencing. MS/MS spectra of immunopeptidome and proteome analysis were searched against a protein database including the human proteome, canonical and non-canonical SARS-CoV-2 ORFs and filtered at 1.5% FDR. **(B)** Population frequency of the 9 endogenous HLA-I alleles expressed in A549 and HEK293T cells. **(C)** Length distribution of HLA peptides in infected and naive cells. **(D)** Motif of 9-mer sequences identified in infected and naive cells. **(E)** Fraction of observed peptides assigned to alleles using HLAthena prediction (percentile rank cutoff <0.5) for immunopeptidome of infected and uninfected cells.

In choosing cell types for this study, we focused on achieving both biological relevance and HLA-I allelic coverage of the human population. A549 are lung carcinoma cells representing the key biological target of SARS-CoV-2, and thus commonly used in COVID-19 studies. HEK293T cells endogenously express HLA-A*02:01 and B*07:02, two high frequency HLA-I alleles (AFA: 12.5%, API:9.5%, HIS:19.4%, EUR: 29.6%, USA: 24.2% for A*02:01; and AFA: 7.3%, API:2.6%, HIS:5.4%, EUR: 14.0%, USA: 10.8% for B*07:02 (Gragert et al., 2013; Poran et al., 2020b)). The nine HLA-I alleles expressed by HEK293T and A549 cells cover 63.8% of the human population (**Fig. 1B**, methods), giving us confidence that our results would be broadly applicable.

Immunopeptidome analysis of high-containment pathogens is technically challenging and requires implementation of protocols that achieve complete virus inactivation while maintaining the integrity of the HLA-peptide complex. We therefore revised our protocol to accomodate the needs and restrictions associated with working in a biosafety level 3 (BSL3) laboratory. First, we used Benzonase nuclease instead of sonication to shear the genomic DNA before immunoprecipitation (Abelin et al., 2019). Second, we inactivated the virus using a lysis buffer containing 1.5% Triton-X, thereby obviating the need to boil or chemically process the cell lysates. Our preliminary investigation showed that these experimental conditions achieved complete virus inactivation (Fig. S1).

We validated the technical performance of our assays by examining the overall characteristics of presented HLA peptides. We identified 5,837 and 6,372 HLA-bound 8-11mer peptides in uninfected and infected A549 cells, and 4,281 and 1,336 unique peptides in HEK293T cells, respectively (Table S1). The reduction in the total number of peptides after infection in HEK293T cells is most likely due to cell death (~50% of cells 24hpi). As expected, we found that the peptide length distribution was not influenced by the virus infection, and the majority of HLA-I peptides were 9-mers (**Fig. 1C**). We focused on 9-mer peptides for subsequent binding motif analyses and compared the peptide amino acid composition between uninfected and infected cells. We did not find major differences in peptide motifs following infection and observed the same amino acids at the main anchor positions 2 and 9 in line with the expected binding motifs of the alleles expressed in the two cell lines (**Fig. 1D**).

To evaluate if the LC-MS/MS-detected peptides are indeed predicted to bind to the expressed HLA-I alleles, we inferred the most likely allele to which each peptide binds using HLAthena predictions (Sarkizova et al., 2020). A549 cells express A*25:01/30:01, B*18:01/44:03 and C*12:03/16:01, while HEK293T cells express A*02:01, B*07:02 and C*07:02 (determined by HLA allotyping). At a stringent cutoff of predicted percentile rank <=0.5, 87% of A549 and 73% of HEK293T identified peptides post infection were assigned to at least one of the alleles in the corresponding cell line (**Fig. 1E**). The majority of presented peptides were assigned to A*25:01 and A*02:01 for A549 and HEK293T cells, respectively (**Fig. 1E**); infection with SARS-CoV-2 did not alter this distribution. Interestingly, a larger fraction (18-24%) of peptides were predicted to bind to multiple alleles including HLA-C alleles in the A549 cell line. HLA-C binding peptides are often deprioritized compared to HLA-A and HLA-B peptides in both epitope prediction and T cell assay studies, yet these data suggest HLA-C peptides should be considered in future analyses.

### Detecting SARS-CoV-2 HLA-I peptides from canonical ORFs

Next, we examined HLA-I peptides that are derived from canonical SARS-CoV-2 proteins (**Fig. 2A**). We identified 15 peptides in A549 cells and 7 in HEK293T cells that are mapped to the viral nsp1, nsp2, nsp3, nsp8, nsp10, nsp14, ORF7a, S, M, and N proteins (RefSeq NC_045512.2). In addition to the use of a stringent FDR cut-off of 1.5% for automated interpretation of peptide spectrum matches from our LC-MS/MS data, SARS-CoV-2-derived peptide identifications were designated high, medium, or low confidence using i) synthetic peptides measurements (for 21 epitopes) and ii) by manually inspecting the spectra (see Methods). Most of the HLA-I peptides were detected in more than one experiment and predicted as good binders by HLAthena to at least one of the expressed HLA alleles. All the tested synthetic peptides validated the viral peptides that we observed in infected cells (Table 1). FASEAARVV from nsp2, IRQEEVQEL from ORF7a, KRVDWTIEY from nsp14, and YLNSTNVTI from nsp3 were also independently confirmed in previously generated biochemical binding assays (Covid19 Intavis_Immunitrack stability dataset 1, https://www.immunitrack.com/free-coronavirus-report-for-download/). One peptide, HADQLTPTW, was also detected in non-infected A549 cells and thus, we removed it from all subsequent analyses.

**Figure 2.**
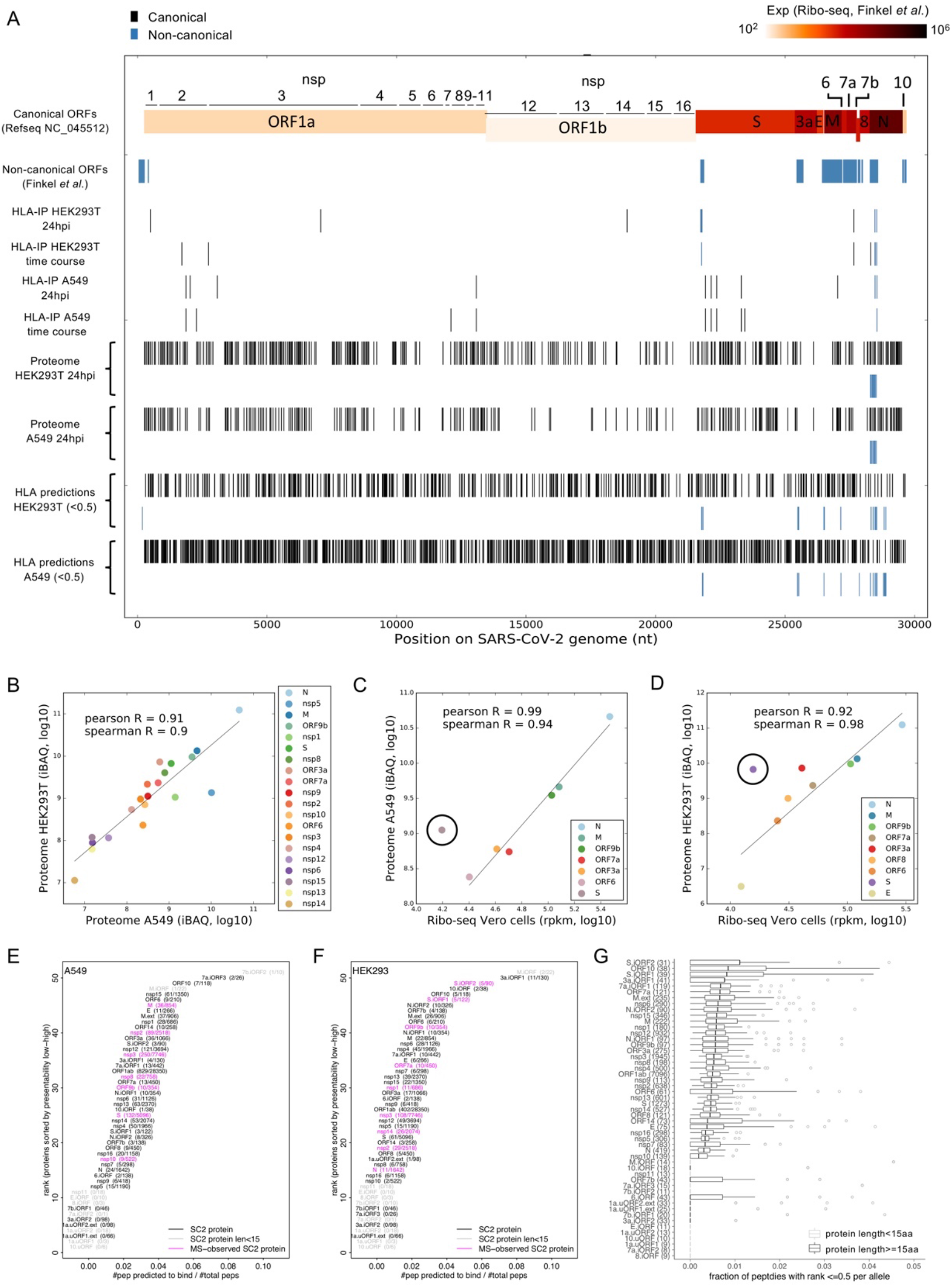
SARS-CoV-2 HLA-I immunopeptidome and whole proteome. **(A)** Summary of peptides location across SARS-CoV-2 genome from HLA-I immunopeptidome, whole proteome, and predictions. Individual tracks represent: (i) Canonical ORFs according to NC_045512.2. Colors represent Ribo-seq translation measurements in Vero cells (Finkel et al., 2020b). (ii) Position of 23 non-canonical ORFs identified by Ribo-seq (Finkel et al., 2020b). (iii) HLA-I peptides detected in HEK293T cells 24hpi. (iv) Total HLA-I peptides detected in a time-course experiment in HEK293T cells 12, 18, and 24hr post infection. (v) HLA-I peptides detected in A549 cells 24hpi. (vi) Total HLA-I peptides detected in a timecourse experiment in A549 cells 12, 18, and 24hr post infection (vii+viii) Tryptic peptides from whole proteome mass-spectrometry in HEK293T and A549 cells. (ix+x) Predicted HLA-I peptides according to HLAthena, rank threshold=0.5. Black and blue bars indicate peptides from canonical and non-canonical ORFs, respectively. **(B)** SARS-CoV-2 proteins abundance in A549 and HEK293T cells 24hpi. iBAQ - intensity-based absolute quantification**. (C, D)** Comparison between our protein abundance measurements 24hpi and ribo-seq (Finkel et al., 2020b) in A549 (C) and HEK293T (D). nsps are not indicated since expression measurements are not comparable between polyproteins (measured by Ribo-seq) and individual cleaved nsps proteins (measured by mass spectrometry). **(E)** SARS-CoV-2 ORFs HLA-I presentation potential in A549 cells. ORFs were ranked according to the ratio between the number of peptides predicted to bind any of the six HLA-I alleles in A549 and the total number of 8-11mers. **(F)** Similar to (E) for HEK293T cells. **(G)** Presentation potential across 92 HLA-I alleles. Showing box plots of the ratio between the number of peptides predicted to bind each of the 92 HLA-I alleles and total number of peptides. ORFs are ranked according to the median across alleles.

**Table 1.**
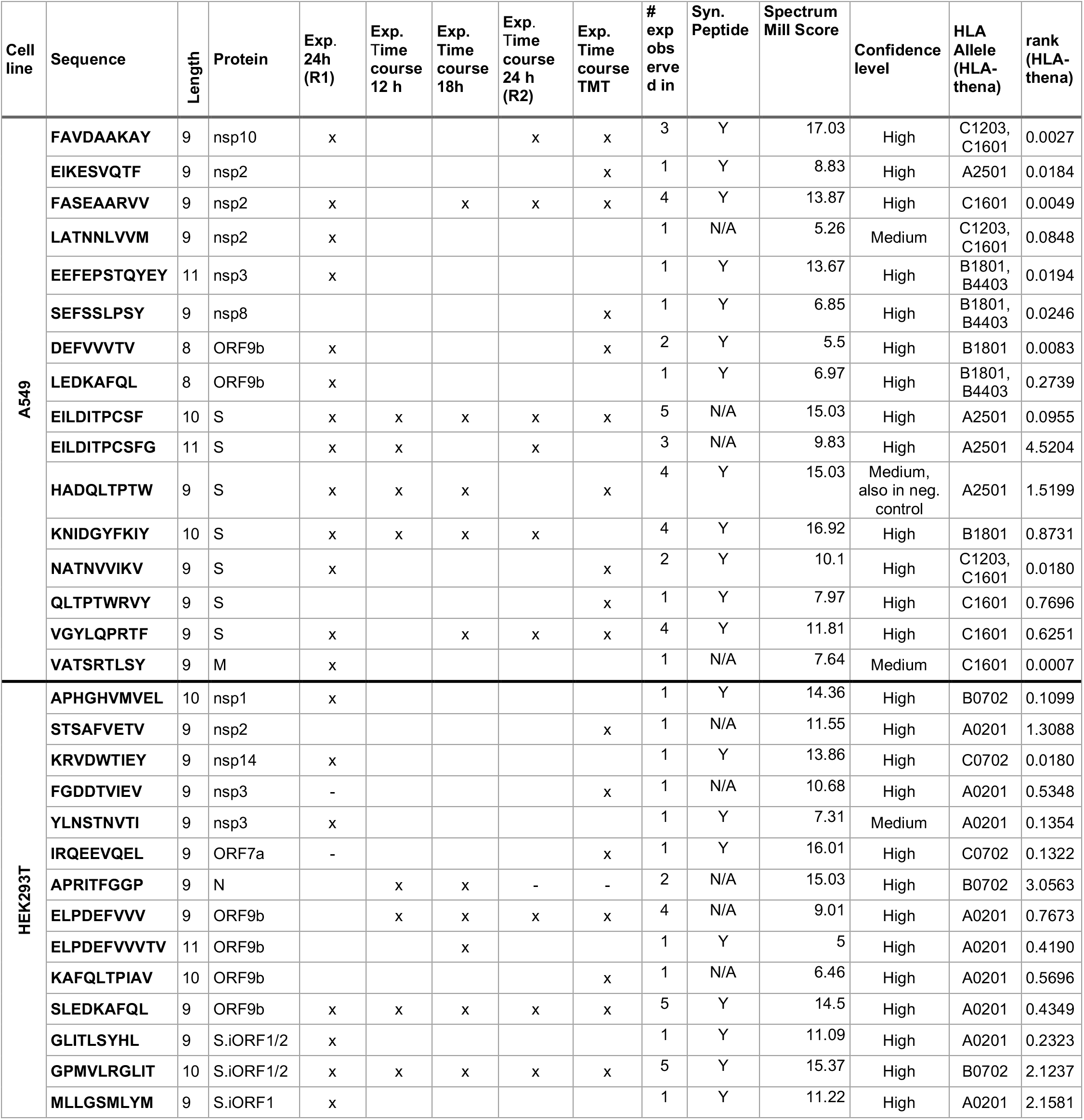
SARS-CoV-2 HLA-I peptides. HLA-I bound peptide sequences from SARS-CoV-2 proteins identified in both A549 and HEK293T cells. Cysteines were detected in their carbamidomethylated or cysteinylated modification state, methionines were oxidized. “x” denotes identification in individual experiments, “-” indicates that the peptide was detected but did not pass FDR correction in that sample, confidence level in identification was evaluated based on Spectrum Mill Score, after manual inspection of MS/MS spectra and comparison to synthetic peptide spectra for available peptides (Y= validated by synthetic peptide, N/A = synthetic peptide not available). For binding prediction using HLAthena, residues were considered unmodified, columns show the most likely HLA allele a peptide is bound to in the respective cell line as well as the percentile binding rank.

Surprisingly, we detect only one HLA-I peptide with an atypical HLA-B*07:02 binding motif from N, a SARS-CoV-2 protein expected to be highly abundant based on previous RNA-seq and Ribo-seq studies (Finkel et al., 2020b; Kim et al., 2020). To test if this low representation could be explained by lower expression of N in our experiment, we examined the whole proteome mass spectrometry data generated from the flowthrough of the HLA-IP. We found a strong correlation between the abundance of viral proteins in the proteome of the two cell lines in our study (Pearson R=0.91 and Spearman R=0.90, **Fig. 2B**), and with recently published Ribo-seq translation measurements in infected Vero cells (Finkel et al., 2020b) (Pearson R=0.99, Spearman R=0.94 for A549 and Pearson R=0.92, Spearman R=0.98 for HEK293T, **Fig. 2C,D**, Table S2A). The N protein remained the most abundant viral protein in both cell lines.

We hypothesized another potential explanation for reduced HLA-I presentation of N derived peptides, that the protein harbours fewer peptides compatible with the HLA binding motifs. To test if the low presentation of peptides from N stems from intrinsic properties of the protein sequence, we computed for each SARS-CoV-2 ORF the ratio between the number of peptides that are predicted to be presented by at least one of the HLA-I alleles in each cell line and the number of total 8-11mers. Notably, N harbors less presentable peptides than most SARS-Co-V-2 proteins in both cell lines (**Fig. 2E, F**, Table S2B). We then expanded our analysis to 92 HLA-I alleles with high population coverage and with immunopeptidome-trained predictors (Sarkizova et al., 2020) (**Fig. 2G**, Table S2B). This analysis implied that N is among the least presentable canonical proteins of SARS-CoV-2, exceeded only by nsp10, nsp11 and ORF7b. Nonetheless, we detected an nsp10 derived HLA-I peptide in A549 cells **(Fig. 2A)**. Together, our results hint that N, which is the focus of T cells immune monitoring in COVID-19 patients, may be a less preferable target in comparison to other SARS-CoV-2 proteins.

Our whole proteome analysis uncovered a number of other important insights. While the translation of ORF1a and 1ab, the source polyproteins of nsps 1-11, is 10-1000 fold lower than structural ORFs (Finkel et al., 2020b), we found that the abundance of some non-structural proteins was comparable to that of structural proteins (e.g nsp1 and nsp8, **Fig. 2B**). This observation may explain why we found HLA-I peptides originating from non-structural proteins. Interestingly, although nsps 1-11 are post-translationally cleaved from the same polyproteins, their expression levels were quite variable (**Fig. 2B**). Nsps 12-15, which are post-translationally cleaved from polyprotein 1ab downstream to the frameshift signal, are indeed expressed at lower levels, as expected. Another interesting observation is that the S protein appeared as an outlier in both cell lines with higher expression in the proteome data compared to the Ribo-seq measurements, which suggests it may undergo positive post-translational regulation.

### SARS-CoV-2 HLA-I peptides are derived from internal out-of-frame ORFs in S and N

Remarkably, we detected nine HLA-I peptides processed from internal out-of-frame ORFs present in the coding region of S and N. In the S.iORF1/2 proteins, we detected three HLA-I peptides: GPMVLRGLIT, GLITLSYHL and MLLGSMLYM in HEK293T cells, all of which are predicted to bind HLA alleles in this cell line (**Fig. 3A**). GLITLSYHL is predicted to bind strongly to A*02:01 (rank 0.23), and GPMVLRGLIT and MLLGSMLYM are predicted to bind weakly to B*07:02 and A*02:01 (rank 2.12 and 2.16, respectively), suggesting the potential for widespread presentation of these non-canonical HLA-I peptides in the population. In addition, we detected six HLA-I peptides from ORF9b, an internal out-of-frame ORF in the coding region of N, in A549 and HEK293T cells (**Fig. 3B**). These HLA-I peptides cover overlapping protein sequences in both cell lines and contain binding motifs compatible with the expressed HLA-I alleles. All six HLA-I peptides were predicted to be highly likely to bind to either A*02:01 in HEK293T cells (SLEDKAFQL, KAFQLTPIAV, ELPDEFVVV, and ELPDEFVVVTV, rank 0.43, 0.57, 0.77 and 0.42, respectively) or B*18:01 in A549 cells (LEDKAFQL and DEFVVVTV, rank 0.27 and 0.0008, respectively). To validate the amino acid sequences of these non-canonical peptides, we compared the tandem mass spectra of synthetic peptides to the experimental spectra and observed high correlation between fragment ions and retention times (+/− 2 minutes, **Fig. 3C**). We also performed 6-frame translation on the SARS-CoV-2 genome and searched our immunopeptidome LC-MS/MS data against those sequences, but found no high-confidence peptide spectrum matches amongst the theoretical non-canonical ORFs whose translation was unsupported by Ribo-seq.

**Figure 3.**
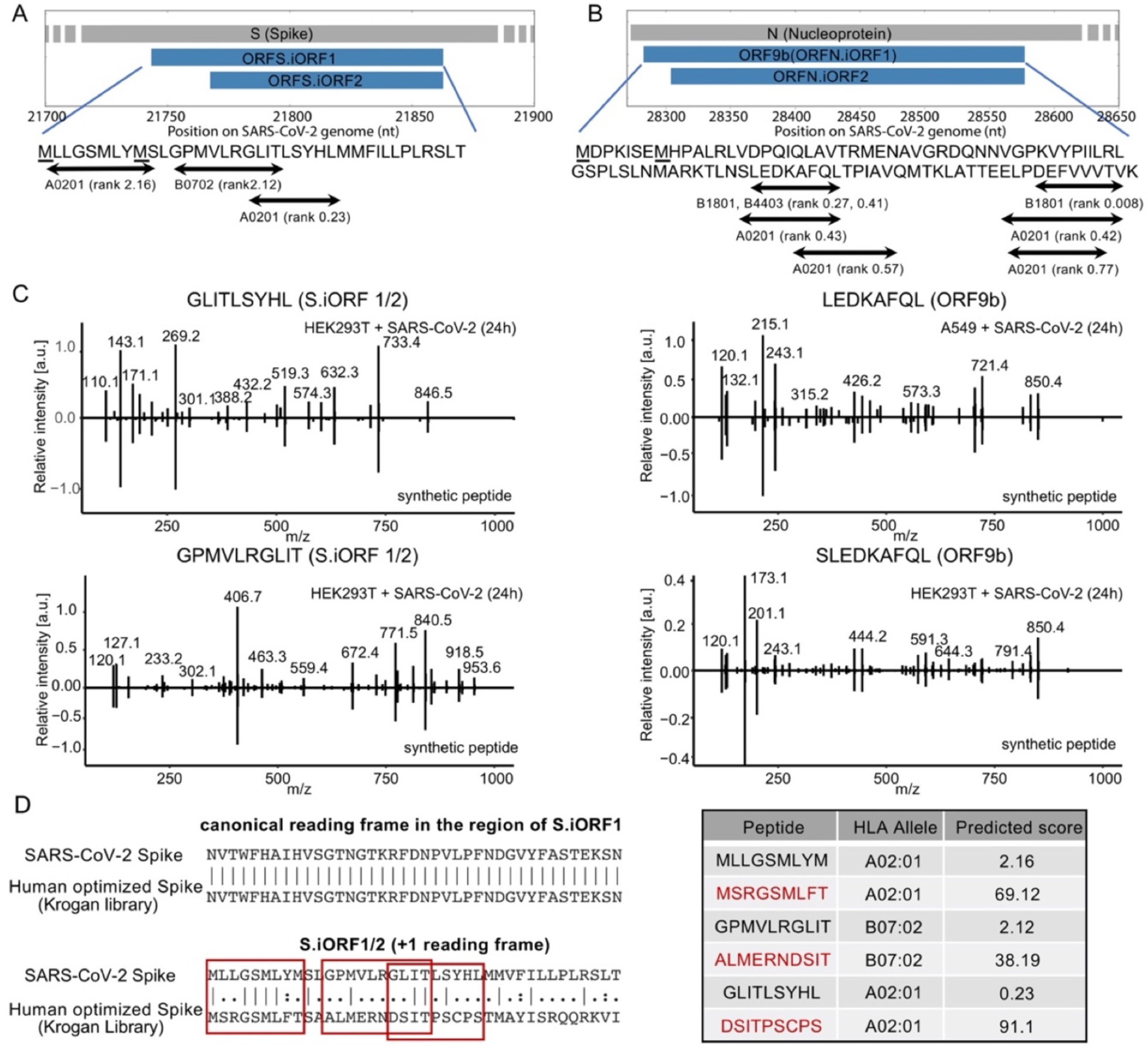
SARS-CoV-2 HLA-I peptides from S.iORF1/2 and ORF9b. **(A)** HLA-I peptides derived from S.iORF1/2. Prediction rank scores were computed with HLAthena. Underscored Methionines represent the start codons of S.iORF1 and S.iORF2. **(B)** HLA-I peptides derived from ORF9b (N.iORF1) and N.iORF2. **(C)** Spectra plots of synthetic peptides confirming the sequences of four HLA-I peptides that were identified in S.iORF1/2 and ORF9b. For each peptide, peaks from HLA-IP samples are presented on the positive y-axis and peaks from synthetic peptides are presented on the negative y-axis. **(D)** The effect of human codon optimization on HLA-I peptides derived from S.iORF1/2. (left) Needleman-Wunsch pairwise global alignment between SASR-CoV-2 sequence and the human optimized S from the Krogan library (Gordon et al., 2020) in the S.iORF1/2 coding region, canonical and +1 reading frames. (right) HLAthena prediction rank scores for the HLA-I peptides detected in our study and the peptides generated by the human optimized S.

In the context of T cell immunity and vaccine development, it is crucial to understand the effect of optimizing RNA sequences on the endogenously processed and presented HLA-I peptides derived from internal out-of-frame ORFs. Two high-profile vaccines under development targeting the S glycoprotein are utilizing RNA-based delivery methods (Callaway, 2020; Jackson et al., 2020; Mulligan et al., 2020) that often involve manipulating the native nucleotide sequence, e.g. via codon optimization, to enhance expression. These techniques maintain the amino acids sequence of the canonical ORF, yet may alter the sequence of proteins encoded in alternative reading frames. We aimed to investigate the effect of optimizing RNA sequences of S on the endogenously processed and presented HLA-I peptides derived from S.iORF1/2. Since the sequences being used in the vaccines are not publicly available, we evaluated the possible effects of codon optimization on HLA-I antigen presentation using the sequence of S from a SARS-CoV-2 human optimized ORFs library (Gordon et al., 2020) in the region encoding S.iORF1/2. As expected, there is no change in the main ORF; however, the amino acid sequence in the +1 frame, which encodes S.iORF1/2, is significantly different (**Fig. 3D**). Since the methionine driving the translation of S.iORF1 is preserved, it is possible that this ORF is expressed in the human optimized construct. However, the peptides likely to be processed from the optimized sequence no longer correspond to native viral peptides and are not predicted to bind the same HLA-I alleles (**Fig. 3D**). These data suggest that RNA-based vaccine design may either preclude the HLA-I presentation of native SARS-CoV-2 peptides derived from overlapping ORFs or promote presentation of alternative peptides. Either case may lead to unintended altering of the immune response.

### HLA-I peptides derived from SARS-CoV-2 are stably presented over the time course of infection

To investigate the dynamics of HLA-I presentation during infection, we performed a time course experiment in which samples for immunopeptidomics, proteomics and RNA-seq were harvested 12, 18 and 24 hours post infection. We chose these time points based on a recent study showing low abundance of SARS-CoV-2 proteins prior to 10 hours post infection (Bojkova et al., 2020). In A549 cells, RNA-seq measurements showed progressively increasing infection, with 47%, 54% and 65% of the cell’s transcriptome mapped to SARS-CoV-2 after 12, 18 and 24 hours, respectively (**Fig. 4A**). In a sharp contrast to the high expression of viral transcripts, only 0.37% of the total protein intensity observed by mass spectrometry was contributed by SARS-CoV-2 at 24h post infection. The low fraction of viral proteins is consistent with the relatively low number of viral HLA-I peptides, 0.43% of the total intensity, observed by mass spectrometry.

**Figure 4.**
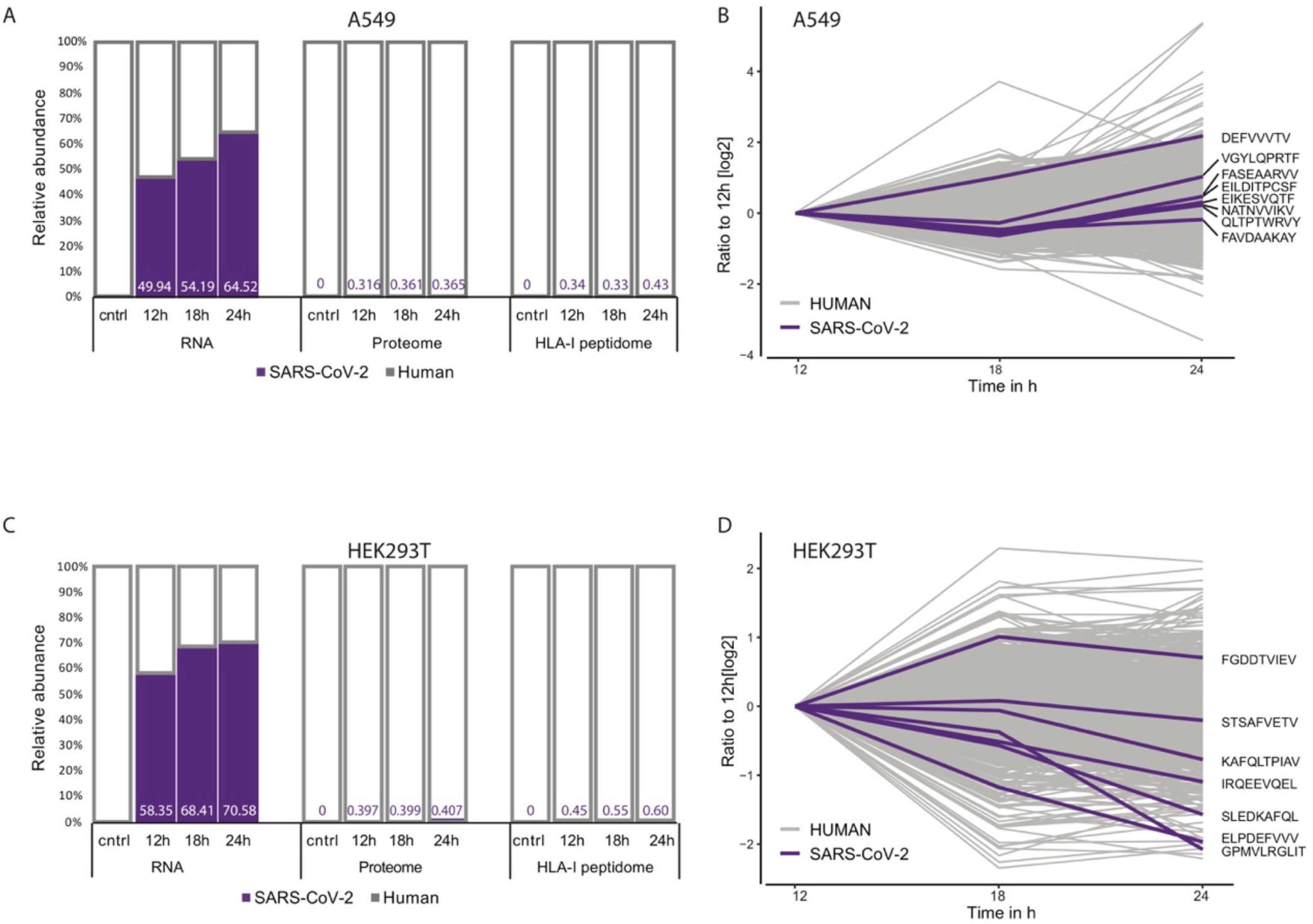
HLA-I peptides dynamics in SARS-CoV-2 infected cells. **(A)** Relative abundance of RNA, proteome and HLA-I peptides derived from SARS-CoV-2 and the human genome in non-infected A549 cells and 12, 18 and 24h post-infection. RNA abundance was calculated as the fraction of reads aligned to SARS-CoV-2 or human transcripts of the total aligned reads. Relative label-free proteome and HLA-I peptidome abundances were determined from the sum of corrected iBAQ values (protein groups) or log-transformed precursor ion chromatographic peak area intensities (HLA peptides) for each precursor ion with a confidently identified peptide in an MS/MS spectrum. **(B)** Dynamics of TMT-labeled HLA-I peptides at the same time points. Intensity values were normalized to the 12h time point. **(C)** Similar to (A), in HEK293T cells **(D)** Similar to (B), in infected HEK293T cells.

To compare the abundance of HLA-I peptides across all three time points we utilized isobaric chemical labeling with tandem mass tags (TMT) for precise relative quantification and to reduce missing values due to stochastic sampling by LC-MS/MS. Labeling with TMT enabled the detection of 8 viral antigens with coverage across all timepoints (**Fig. 4B**, Table 1, Table S1). We observed only a small increase in the abundance of HLA-I peptides over time, except for the ORF9b peptide, DEFVVVTV, that has a log2 ratio of 2.18 24h post infection. Similar trends were observed for human-derived HLA-I peptides, with most of them recording no or only slight increase over time.

We found similar patterns of highly abundant viral transcripts, with overall low representation at the proteome and HLA-I peptidome in HEK293T cells. RNA-seq measurements showed 58%, 68% and 71% of the cell’s transcriptome mapped to SARS-CoV-2 genome after 12, 18 and 24 hours, respectively (**Fig. 4C**). Similarly to A549 cells, only 0.41% of the total proteome and 0.6% of the total HLA-I peptides originated from viral proteins 24h post-infection. We detected six peptides from SARS-CoV-2 with coverage across all three time points in a TMT labeled immunopeptidome experiment (**Fig. 4D**, Table 1, Table S1). Relative peptide abundance decreased for all peptides at the 24h time point compared to 18h, which is likely attributed to the increasing number of dying cells after 24 hours (~50% of the cells in the culture). Overall, we find that SARS-CoV-2 HLA-I peptides are stable over three time points in two independent cell lines.

### SARS-CoV-2 infection interferes with cellular pathways that may impact HLA-I peptide processing and presentation and immune signaling

To assess if SARS-CoV-2 infection alters the HLA-I antigen processing and presentation pathway, we first compared the overlap between HLA-I peptidomes of uninfected and 24hr post-infection cells and found that 62% of these peptides were detected in both experiments (**Fig. 5A**). This degree of overlap is similar to biological replicates of the same sample (Abelin et al., 2017; Demmers et al., 2019; Sarkizova et al., 2020). We then compared the calculated abundance of individual viral proteins and viral HLA-I peptides relative to the human proteome of the cell lines 24h post infection. Although the overall viral abundance in the cell proteome is relatively low, individual viral proteins are highly expressed and exceed most of the cellular proteins (Wilcoxon rank-sum test, p<10^−4^ and p <10^−6^ for A549 and HEK293T cells, respectively, **Fig. 5B** and Fig. S2A, Table S3). In contrast to the high expression of their source proteins, the intensity of viral HLA-I peptides is similar to peptides from the host proteome, indicating that viral peptides are not preferentially presented in infected cells (Wilcoxon rank-sum test, p>0.8 and p>0.4 for A549 and HEK293T cells, respectively, **Fig. 5C**, Fig. S2B, Table S3).

**Figure 5.**
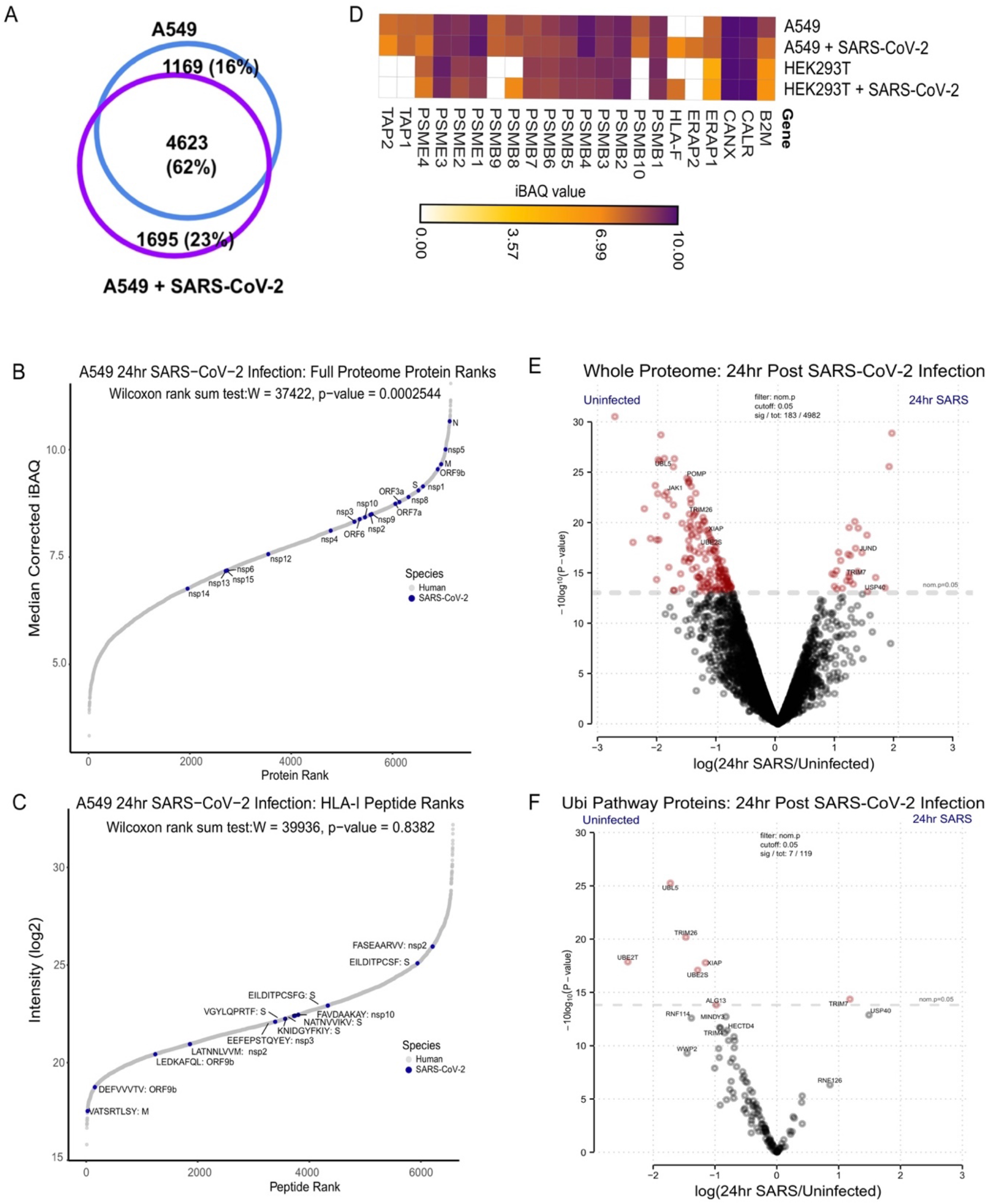
The effect of SARS-CoV-2 infection on antigen presentation in host cells. **(A)** Venn-diagram showing the overlap between total HLA-I peptides in uninfected (blue) and 24 hours post SARS-CoV-2 infection (purple) A549 cells. **(B)** Rank plot of the iBAQ value for each human (gray) and SARS-CoV-2 (blue) protein detected in A549 cells 24 h.p.i. SARS-CoV2 proteins are annotated with their respective gene names. **(C)** Similar rank plot to (B) but for observed HLA-I peptides and their intensities. Peptides mapped to SARS-CoV-2 are annotated with their respective amino acid sequence and source protein gene name. **(D)** Heatmap of iBAQ values for canonical antigen presentation pathway proteins observed across uninfected and infected A549 and HEK293T cells 24 h.p.i **(E)** Volcano plot comparing protein levels across uninfected and infected A549 and HEK293T cells 24 h.p.i. Significantly changing proteins (p-value <0.5; red) are shown along with annotations of specific proteins involved in proteasomal degradation and immune response. **(F)** Similar to (E), a volcano plot for a subset of proteins with functions related to the ubiquitin-proteasome system.

The high overlap of host HLA-I peptides in infected and uninfected cells and the overall low percentage of SARS-CoV-2 in the immunopeptidome, led us to interrogate the whole proteome data for evidence of viral interference with the antigen presentation and protein degradation pathways. Because HLA-A, -B and -C are depleted by the HLA-IP these proteins could not be analyzed in the whole proteome flowthrough. However, all other proteins should remain intact and can be used for proteomic analyses of host responses to infection. We initially evaluated canonical HLA-I presentation pathway proteins across both the uninfected and 24 hr post-infection A549 and HEK293T cells (**Fig. 5D**, Fig. S2C, Table S3) and observed no significant differences in these proteins, such as B2M, ERAP1/2, TAP1/2, and proteasome subunits. Of note, HLA-F, which has been shown to not be reactive to W6/32 antibody used for the IP(Wainwright et al., 2000), and thus should not be depleted from the lysate prior to the whole proteome analysis, showed increased expression upon infection. HLA-F has recently been recognized as an immune regulatory molecule that interacts with KIR3DS1 on NK cells during viral infection (Lunemann et al., 2018). We then compared the entire set of proteins detected across A549 and HEK293T experiments to determine if upstream antigen processing proteins were altered (**Fig. 5E**, Table S3), and found that POMP (Proteasome Maturation Protein), a chaperone critical for the assembly of 20S proteasomes and immunoproteasomes, was significantly depleted in both infected cell lines. POMP has recently been reported to impact ORF9c stability, which has been implicated in suppressing the antiviral response in cells (Dominguez Andres et al., 2020). Several ubiquitination pathway proteins were significantly altered in response to SARS-CoV-2 infection (**Fig. 5F**, Table S3), including depletion of TRIM26, an E3 ligase that modulates IRF3, NF-κB activation and IFN-β induction(Ran et al., 2016), and enrichment of TRIM7, which has recently been shown to bind M (Stukalov et al., 2020). Interestingly, the tyrosine kinase, JAK1, critical for IFN signaling that has been reported to be reduced with SARS-CoV nsp1 overexpression (Wathelet et al., 2007) was depleted in both cell lines upon SARS-CoV-2 infection (**Fig. 5E**). Taken together, these data suggest that SARS-CoV-2 interferes with IFN signaling proteins and the HLA-I pathway through both POMP depletion and by altering ubiquitination enzymes, that in turn, may prevent highly expressed SARS proteins from being processed and presented.

### LC-MS/MS-identified SARS-CoV-2 HLA class I peptides have high predicted population coverage

Increasingly accurate HLA-I presentation prediction tools are routinely applied to the full transcriptome or proteome of an organism to computationally nominate presentable epitopes. However, these tools are trained on data that is agnostic to virus-specific processes that may interfere with the presentation pathway. Thus, the sensitivity and specificity of *in silico* predictions for any particular virus are insufficiently characterized. To assess how well computational tools would recover the HLA-I peptides that we identified by LC-MS/MS, we used HLAthena (predictor trained exclusively on endogenous HLA epitopes)(Abelin et al., 2017; Sarkizova et al., 2020) to retrospectively predict all 8-11mer peptides tiling known SARS-CoV-2 proteins against the complement of HLA-I alleles expressed by A549 and HEK293T (**Fig. 6A**, Table S4A). Of the 29 MS-identified peptides, 18 had a predicted percentile rank <0.5 and 25 had percentile rank <2. The single A549 peptides with %rank >1, EILDITPCSFG, was also observed in a shorter form, EILDITPCSF, with a much more favorable prediction (%rank=0.095), suggesting that a phenylalanine C-terminal anchor is preferable and thus the 11mer could be bound with an overhang. Similarly, GPMVLRGLIT was more favorably predicted as a 9-mer, GPMVLRGLI, although the short version was not detected.

**Figure 6.**
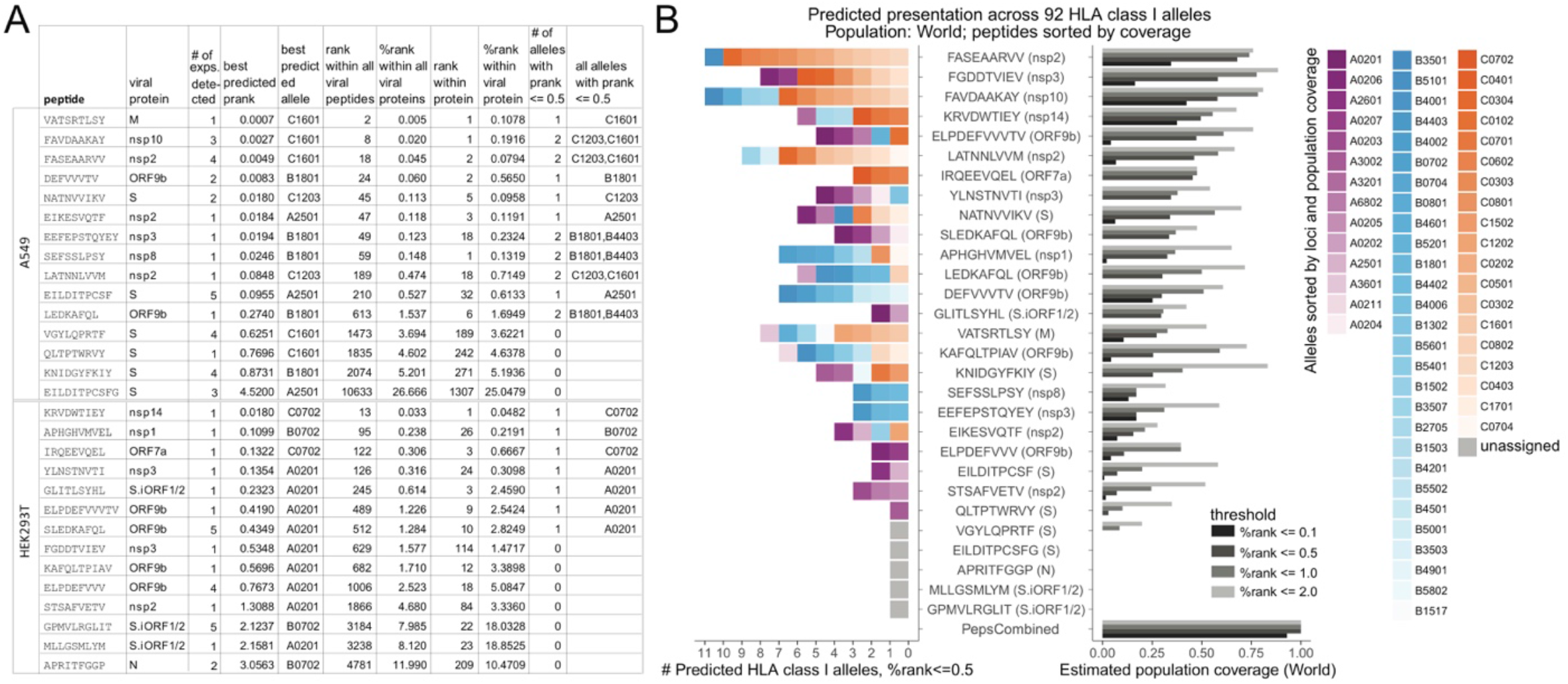
Presentation prediction and population coverage estimates of MS-identified SARS-CoV-2 HLA-I peptides. **(A)** Summary of LC-MS/MS identified SARS-CoV-2 epitopes with corresponding HLAthena predictions for the 6 HLA-I alleles expressed by A549 (HLA-A*25:01, A*30:01, B*18:01, B*44:03, C*12:03, C*16:01; top) and the 3 HLA-I alleles in HEK293T (HLA-A*02:01, B*07:02, C*07:02; bottom). **(B)** HLAthena predictions for 92 HLA-I alleles using percentile rank cutoff values of 0.1, 0.5, 1, and 2%, demonstrating the number of unique HLA-A, -B, and -C alleles that LC-MS/MS observed SARS-CoV-2 peptides are predicted to strongly bind (left) to achieve a high estimated population coverage (right). Alleles are colored and ordered according to loci and world population frequency (high to low color intensity). Peptides are ordered according to their estimated coverage at %rank cutoff of 0.5.

Within a total of 39,875 SARS-CoV-2 8-11mers, 11 out of 15 A549 peptides and 9 out of 14 HEK293T peptides had percentile rank scores within the top 1000 viral peptides (top 1.5% and 1.7% for A549 and HEK293T, respectively). To account for variability in viral protein expression levels, we also performed this analysis within the source protein of each peptide. We found that 13 out of 29 peptides scored within the top 10 epitopes amongst all 8-11mers that the source protein can give rise to, and 17 scored within the top 20. Overall, predicted ranks were higher for A549 compared to HEK293T, which could partially be due to the larger number of viable cells and deeper peptidome we obtained for A549. These observations suggest that while an *in silico* epitope prediction scheme that nominates the top 10-20 peptides of each viral protein would recover ~50% (13-17 of 29) of observed epitopes with very high priority, this short list would still only encompass ~5-10% true LC-MS/MS positives.

Next, we estimated the HLA allele coverage achieved by the observed endogenously processed and presented viral epitopes amongst AFA, API, EUR, HIS, USA, and World populations at different rank cutoffs (top 0.1%, 0.5%, 1% and 2%) based on HLAthena predictions across 92 HLA-A, -B, and -C alleles (**Fig. 6B**, Fig. S3, Table S4B,C). At the second most stringent cutoff, 0.5%, 24 of the 29 individual peptides were predicted to bind at least one allele (range: 1-11, median: 5, mean: 5.3) with 3-68% (median: 30%, mean: 31%) world coverage. Combined together, the MS-identified peptide pool was estimated to cover at least one HLA-A, -B, or -C allele for 99% of the population with at least one peptide. These estimates indicate that HLA-I immunopeptidomics on only two cell lines combined with accurate epitope prediction tools can rapidly prioritize multiple CD8+ T cell epitopes with high population coverage for COVID-19 immune monitoring and vaccine development.

## DISCUSSION

We provide the first view of SARS-CoV-2 HLA-I peptides that are endogenously processed and presented by infected cells. As expected, we observed thousands of distinct HLA-I peptides derived mostly from self proteins(Schellens et al., 2015a, 2015b). We detected almost half of the virus-derived peptides in independent experiments and in different time points. Remarkably, 9 of 29 viral peptides detected are derived from internal out-of-frame ORFs in S (S.iORF1/2) and N (ORF9b). This observation implies that current studies on T cell responses in COVID-19 patients that focus on the canonical viral ORFs, like those that leverage MegaPool epitopes(Grifoni et al., 2020a; Weiskopf et al., 2020), exclude a significant number of virus-derived HLA-I epitopes. Moreover, many of these studies do not include peptides that are predicted to bind HLA-C, a classical HLA that may not have been considered due to its lower expression compared to HLA-A and -B. However, 9 (~30%) of the HLA-I peptides detected in our experiment, including peptides derived from S, are predicted as good binders to HLA-C alleles. Together, these findings hint that current peptide pools may be incomplete. Therefore, unbiased approaches such as LC-MS/MS immunopeptidomics that directly identify naturally presented SARS-CoV-2 peptides on host cells can inform the design of future pools to allow more comprehensive characterization of T cell responses in COVID-19 patients.

Our analyses demonstrate that synthetic approaches aiming at enhancing the expression of canonical ORFs, some of which may be utilized in current vaccine strategies, can inadvertently eliminate or alter the production of HLA-I peptides derived from overlapping reading frames. The large number of HLA-I peptides detected from internal out-of-frame ORFs suggests that accounting for these ORFs may considerably contribute to the induction of effective memory T cells in response to SARS-CoV-2 vaccines that overexpress S. Researchers may need to carefully examine the effects of sequence manipulation and codon optimization on the amino acids encoded by S.iORF1 and S.iORF2 with emphasis on the three HLA-I peptides that we identified experimentally: GPMVLRGLIT, GLITLSYHL and MLLGSMLYM. In broader terms, many viral genomes have evolved to increase their coding capacity by utilizing overlapping ORFs and programmed frameshifting(Ketteler, 2012). Thus, our data emphasise the importance of combining unbiased approaches to detect the complete translatome and immunopeptidome of viruses in order to design vaccines with optimal antibody and T cells responses.

Proteomics analyses of infected cells show that SARS-CoV-2 may interfere with the presentation of HLA-I peptides and the expression of ubiquitination and immune signaling pathway proteins. We found that SARS-CoV-2 infection leads to a significant decrease in POMP expression and alters the expression of ubiquitination pathway proteins. By impacting ubiquitin mediated proteasomal degradation and immune signaling proteins, SARS-CoV-2 may reduce the precursors for downstream processing and HLA-I presentation and alter the immune response. Furthermore, IFN signaling protein JAK1 was depleted in both cell lines post infection, which could relate to the delayed IFN response observed in COVID-19 patients (Acharya et al., 2020). Although follow-up is required to determine the mechanistic role of JAK1 in SARS-CoV-2 infection, these data provide candidate proteins that can be investigated to uncover how SARS-CoV-2 impacts HLA-I antigen presentation and proteins involved in immune signaling pathways.

Surprisingly, we detect only one atypical HLA-I peptide derived from N. Using epitope predictions from 92 frequent HLA-A, -B, and -C alleles, we show that N is less likely to generate presentable peptides in comparison to other ORFs of SARS-CoV-2. An intriguing hypothesis is that viruses have evolved to attenuate HLA-I presentation of highly abundant proteins as an immune evasion mechanism. Yet, SARS-CoV-2 has only recently spilled over to humans from bats, so human HLA-I could not play a role in its evolution. Studying HLA-I presentation of SARS-CoV-2 in bats as well as expanding the relative presentability analysis to other human Coronaviruses are needed in order to evaluate the selective pressure shaping the evolution of N and whether it was adapted to attenuate HLA-I presentation. It is plausible that lacking peptides from N with typical binding motifs represents a sensitivity limitation of our method, the differences of antigen processing machinery within our cell lines or the time points post infection, as earlier or later time points may reveal additional HLA-I epitopes. Recent studies in COVID-19 patients indeed detected positive CD8+ T cell responses against pooled peptides from N (Grifoni et al., 2020a; Le Bert et al., 2020). However, T cell reactivity toward N was lower than S, M and nsp6 (accounting for 12%, 26%, 22% and 15% of the reactivity, respectively)(Grifoni et al., 2020a) even though their source protein abundance is at least 10-fold lower (**Fig. 2B**). Together with these clinical observations, our findings support the notion that N may be less immunogenic than expected based on its high expression level, and that ORF9b and nonstructural proteins should be considered in multiepitope vaccine pools and immune monitoring efforts that currently dominated by highly abundant structural proteins like N and M.

Although only nine HLA-I alleles were profiled in our study, we show that these data combined with *in silico* epitope prediction can rapidly prioritize peptides that cover a wide range of HLA-I alleles. Importantly, two of the HLA-I alleles, A*02:01 and B*07:02, have high frequencies across subpopulations and are prioritized for *in-vitro* and *in-vivo* studies (Ferretti et al.; Grifoni et al., 2020a), allowing a direct comparison between our results and ongoing efforts. We found that most LC-MS/MS-detected HLA-I viral epitopes have very strong predicted binding scores with 9 of 29 ranked within the top 3 source protein peptides. Nevertheless, prediction models are not equipped to explain why viral peptides with comparably strong predicted binding, especially from the same source protein, are not detected, where the lack of detection should not simply be attributed to MS sensitivity since nearly half of the viral peptides are detected in multiple experiments. Epitope prediction models do not incorporate virus-specific processing and presentation preferences that can be ascertained by sequencing HLA-I peptides from SARS-CoV-2 infected host cells. Future studies of SARS-CoV-2 infected cell lines from diverse lineages and primary tissues will likely facilitate virus-specific processing and presentation rules that can be integrated into epitope prediction models to improve accuracy. Furthermore, combining epitope prediction methods and LC-MS/MS to investigate HLA-II peptide presentation in SARS-CoV-2 infected cells may also reveal novel epitopes and epitope presentation rules, as the HLA-II pathway distinctly involves both endocytosis and autophagy, and functions in specific antigen presenting cell populations. HLA-II epitope profiling efforts will also aid in our understanding of both T cell and B cell responses.

In conclusion, the discovery of endogenously presented viral HLA-I peptides and biological insights into how SARS-CoV-2 may alter antigen presentation pathways will enable a more precise selection of peptides for COVID-19 immune monitoring and vaccine development.

## ACKNOWLEDGEMENTS

This project was started in April 2020 while SARS-CoV-2 was fast spreading in the US and the people of Boston were sheltering in place. We are grateful to the tremendous efforts of the supporting teams, both at the Broad Institute and the NEIDL, who facilitated our research under those challenging circumstances. We thank Orel Mizrahi, Suzanna Rachimi and Erica Normandin for technical assistance, DR Mani and Hayden Metsky for computational assistance and Tamara Ouspenskaia and Pierre M. Jean-Beltran for insightful discussions. This study was supported in part by grants from the National Institute of Allergy and Infectious Diseases (U19AI110818 to P.C.S) and the National Cancer Institute (NCI) Clinical Proteomic Tumor Analysis Consortium grants (NIH/NCI U24-CA210986 and NIH/NCI U01 CA214125 to S.A.C and NIH/NCI U24CA210979 to D.R.M). S.W-G. is the recipient of the Human Frontier Science Program fellowship (LT-000396/2018), EMBO non stipendiary Long-Term Fellowship (ALTF 883-2017), the Gruss-Lipper Postdoctoral Fellowship, the Zuckerman STEM Leadership Program Fellowship and the Rothschild Postdoctoral Fellowship. S.K is a Cancer Research Institute/Hearst foundation fellow. C.H.T.T. was supported by a National Science Foundation Graduate Research Fellowship (Grant No. 1745303). M.G. is the recipient of an EMBO Long-Term Fellowship (ALTF 486-2018) and a Cancer Research Institute/Bristol-Myers Squibb Fellow (CRI2993). D.B.K acknowledges funding support from Emerson Collective and NIH/NCI R21 CA216772-01A1. C.M.R is supported by G. Harold and Leila Y. Mathers Charitable Foundation, the Bawd Foundation, and NIH NIAID 3 R01-AI091707-10S1. M.S. is supported by Boston University startup funds.

## AUTHOR CONTRIBUTIONS

S.W-G., S.K., S.S. and J.G.A. conceptualized the study. S.W-G., S.K., J.G.A, C.M.R and M.S. designed the experiments. S.K., S.S., L.R.P., D.C., M.R.B., H.B.T, H.L.C, K.R., D.B.K., J.G.A, M.S., performed experiments, S.W-G., S.K., S.S., M.R.B., C.H.T-T, Y.F., A.N., M.G., K.R.C., J.G.A. performed data analysis. S.W-G., S.K., S.S., K.R.C, N.H., S.A.C, J.G.A, M.S., and P.C.S wrote the manuscript with comments from all the authors.

## METHODS

### Cell culture

Human embryonic kidney HEK293T cells, human lung A549 cells, and African green monkey kidney Vero E6 cells were maintained at 37°C and 5% CO2 in DMEM containing 10% FBS. We generated stable HEK293T and A549 cells expressing human ACE2 and TMPRSS2 by transducing them with lentivirus particles carrying these two cDNAs.

### SARS-CoV-2 virus stock preparation

The 2019-nCoV/USA-WA1/2020 isolate (NCBI accession number: MN985325) of SARS-CoV-2 was obtained from the Centers for Disease Control and Prevention and BEI Resources. To generate the virus P1 stock, we infected Vero E6 cells with this isolate for 1h at 37°C, removed the virus inoculum, rinsed the cell monolayer with 1X PBS, and added DMEM supplemented with 2% FBS. Three days later, when the cytopathic effect of the virus became visible, we harvested the culture medium, passed through a 0.2μ filter, and stored it at −80°C. To generate the virus P2 stock, we infected Vero E6 cells with the P1 stock at a multiplicity of infection (MOI) of 0.1 plaque forming units (PFU)/cell and harvested the culture medium three days later using the same protocol as for the P1 stock. All experiments in this study were performed using the P2 stock.

### Plaque assay

Vero E6 cells were used to determine the titer of our virus stock and to evaluate SARS-CoV-2 inactivation following lysis of infected cells in our HLA-IP buffer. Briefly, we seeded Vero E6 cells into a 12-well plate at a density of 2.5 × 10^5^ cells per well, and the next day, infected them with serial 10-fold dilutions of the virus stock (for titration) or the A549 lysates (for the inactivation assay) for 1h at 37°C. We then added 1 ml per well of the overlay medium containing 2X DMEM (Gibco: #12800017) supplemented with 4% FBS and mixed at a 1:1 ratio with 1.2% Avicel (DuPont; RC-581) to obtain the final concentrations of 2% and 0.6% for FBS and Avicel, respectively. Three days later, we removed the overlay medium, rinsed the cell monolayer with 1X PBS and fixed the cells with 4% paraformaldehyde for 30 minutes at room temperature. 0.1% crystal violet was used to visualize the plaques.

### Immunoprecipitation of HLA-I peptide complexes

Cells engineered to express SARS-CoV-2 entry factors were seeded into nine 15 cm dishes (three dishes per time point) at a density of 15 million cells per dish for A549 cells and 20 million cells per dish for HEK293T cells. The next day, the cells were infected with SARS-CoV-2 at a multiplicity of infection (MOI) of 3. To synchronize infection, the virus was bound to target cells in a small volume of opti-MEM on ice for one hour, followed by addition of DMEM/2% FBS and switching to 37°C. At 12, 18, and 24h postinfection, the cells from three dishes were scraped into 2.5ml/dish of cold lysis buffer (20mM Tris, pH 8.0, 100mM NaCl, 6mM MgCl2, 1mM EDTA, 60mM Octyl β-d-glucopyranoside, 0.2mM Iodoacetamide, 1.5% Triton X-100, 50xC0mplete Protease Inhibitor Tablet-EDTA free and PMSF) obtaining a total of 9 ml lysate. This lysate was split into 6 eppendorf tubes, with each tube receiving 1.5 ml volume, and incubated on ice for 15 min with 1ul of Benzonase (Thomas Scientific, E1014-25KU) to degrade nucleic acid. The lysates were then centrifuged at 4,000 rpm for 22min at 4°C and the supernatants were transferred to another set of six eppendorf tubes containing a mixture of pre-washed beads (Millipore Sigma, GE17-0886-01) and 50 ul of an MHC class I antibody (W6/32) (Santa Cruz Biotechnology, sc-32235). The immune complexes were captured on the beads by incubating on a rotor at 4°C for 3hr. The unbound lysates were kept for whole proteomics analysis while the beads were washed to remove non-specifically bound material. In total, nine washing steps were performed; one wash with 1 mL of cold lysis wash buffer (20mM Tris, pH 8.0, 100mM NaCl, 6mM MgCl2, 1mM EDTA, 60mM Octyl β-d-glucopyranoside, 0.2mM Iodoacetamide, 1.5% Triton X-100), four washes with 1mL of cold complete wash buffer (20mM Tris, pH 8.0, 100mM NaCl, 1mM EDTA, 60mM Octyl β-d-glucopyranoside, 0.2mM Iodoacetamide), and four washes with 20mM Tris pH 8.0 buffer. Dry beads were stored at −80°C until mass-spectrometry analysis was performed.

### HLA-I immunopeptidome LC-MS/MS data generation

HLA peptides were eluted and desalted from beads as described previously(Sarkizova et al., 2020). After the primary elution step, HLA peptides were reconstituted in 3% ACN/5% FA and subjected to microscaled basic reverse phase separation. Briefly, peptides were loaded on Stage-tips with 2 punches of SDB-XC material (Empore 3M) and eluted in three fractions with increasing concentrations of ACN (5%, 10% and 30% in 0.1% NH4OH). For the time course experiment, one third of each sample was also labeled with TMT6 (Thermo Fisher Scientific, lot × UC280588, A549: 12h:126, 18h:128, 24h:130, HEK293T: 12h:127, 18h:129, 24h:131)(Thompson et al., 2003), combined and desalted on C18, and then eluted with increasing concentrations of ACN (10%, 15% and 50%) in 5 mM Ammonium formate. Peptides were reconstituted in 3% ACN/5%FA prior to loading onto an analytical column (25-30cm, 1.9um C18 (Dr. Maisch HPLC GmbH), packed in-house PicoFrit 75 μm inner diameter, 10 μm emitter (New Objective)). Peptides were eluted with a linear gradient (EasyNanoLC 1200, Thermo Fisher Scientific) ranging from 6-30% Solvent B (0.1%FA in 90% ACN) over 84 min, 30-90% B over 0 min and held at 90% B for 5 min at 200 nl.min. MS/MS were acquired on a Thermo Orbitrap Exploris 480 equipped with FAIMS (Thermo Fisher Scientific) in data dependent acquisition. FAIMS CVs were set to −50 and - 70 with a cycle time of 1.5s per FAIMS experiment. MS2 fill time was set to 100ms, collision energy was 29CE or 32CE for TMT respectively.

### Whole proteome LC-MS/MS data generation

200 uL aliquot of HLA IP supernatants were reduced for 30 minutes with 5mM DTT (Pierce DTT: A39255) and alkylated with 10mM IAA (Sigma IAA: A3221-10VL) for 45 minutes both at 25°C on a shaker (1000 rpm). Protein precipitation using methanol/chloroform was then performed. Briefly, methanol was added at a volume of 4x that of HLA IP supernant aliquot. This was followed by a 1x volume of chloroform and 3x volume of water. The sample was mixed by vortexing and incubated at −20° C for 1.5 hours. Samples were then centrifuged at 14,000 rpm for 10 minutes and the upper liquid layer was removed leaving a protein pellet. The pellet was rinsed with 3x volume of methanol, vortexed lightly, and centrifuged at 14,000 rpm for 10 minutes. Supernatant was removed and discarded without disturbing the pellet. Pellets were resuspended in 100 mM triethylammonium bicarbonate (pH 8.5) (TEAB). Samples were digested with LysC (1:50) for 2 hr on a shaker (1000 rpm) at 25°C, followed by trypsin (1:50) overnight. Samples were acidified by 1% formic acid final concentration and dried. Samples were reconstituted 4.5 mM ammonium formate (pH 10) in 2% (vol/vol) acetonitrile and separated into four fractions using basic reversed phase fractionation on a stagetip. Fractions were eluted at 5%, 12.5%, 15%, and 50% ACN/4.5 mM ammonium formate (pH 10) and dried. Fractions were reconstituted in 3%ACN/5%FA, and 1 ug was used for LC-MS/MS analysis. MS/MS were acquired on a Thermo Orbitrap Exploris 480 (Thermo Fisher Scientific) in data dependent acquisition with FAIMS (MS2 isolation width 0.7m/z, cycle time 0.8, collision energy 30% for each CV) and without (MS2 isolation width 0.7m/z, top20 scans, collision energy 30%). Uninfected 1 ug single shot samples were analyzed similarly. For the time course experiment, the samples (12h, 18h, 24h) were not fractionated and 1 ug was used for LC-MS/MS analysis, as described above except that FAIMS with −50, −65, and −85 CV was applied.

### LC-MS/MS data interpretation

Peptide sequences were interpreted from MS/MS spectra using Spectrum Mill (v 7.1 pre-release) to search against a RefSeq-based sequence database containing 41,457 proteins mapped to the human reference genome (hg38) obtained via the UCSC Table Browser (https://genome.ucsc.edu/cgi-bin/hgTables) on June 29, 2018, with the addition of 13 proteins encoded in the human mitochondrial genome, 264 common laboratory contaminant proteins, 553 human non-canonical small open reading frames, 28 SARS-CoV2 proteins obtained from RefSeq derived from the original Wuhan-Hu-1 China isolate NC_045512.2 (https://www.ncbi.nlm.nih.gov/nuccore/1798174254)(Wu et al., 2020), and 23 novel unannotated virus ORFs whose translation is supported by Ribo-seq (Finkel et al., 2020b) for a total of 42,337 proteins. Amongst the 28 annotated SARS-CoV2 proteins we opted to omit the full-length polyproteins ORF1a and ORF1ab, to simplify peptide-to-protein assignment, and instead represented ORF1ab as the mature 16 individual non-structural proteins that result from proteolytic processing of the 1a and 1ab polyproteins. We further added the D614G variant of the SARS-Cov2 Spike protein that is commonly observed in European and American virus isolates.

For immunopeptidome data MS/MS spectra were excluded from searching if they did not have a precursor MH+ in the range of 600-4000, had a precursor charge >5, or had a minimum of <5 detected peaks. Merging of similar spectra with the same precursor m/z acquired in the same chromatographic peak was disabled. Prior to searches, all MS/MS spectra had to pass the spectral quality filter with a sequence tag length >1 (*i.e*., minimum of 3 masses separated by the in-chain masses of 2 amino acids) based on HLA v3 peak detection. MS/MS search parameters included: ESI-QEXACTIVE-HCD-HLA-v3 scoring parameters; no-enzyme specificity; fixed modification: carbamidomethylation of cysteine; variable modifications: cysteinylation of cysteine, oxidation of methionine, deamidation of asparagine, acetylation of protein N-termini, and pyroglutamic acid at peptide N-terminal glutamine; precursor mass shift range of −18 to 81 Da; precursor mass tolerance of ±10 ppm; product mass tolerance of ± 10 ppm, and a minimum matched peak intensity of 30%. Peptide spectrum matches (PSMs) for individual spectra were automatically designated as confidently assigned using the Spectrum Mill auto-validation module to apply target-decoy based FDR estimation at the PSM level of <1.5% FDR. For the TMT-labeled time course experiments, two parameters were revised: the MH+ range filter was 800-6000, and TMT labeling was required at lysine, but peptide N-termini were allowed to be either labeled or unlabeled. Relative abundances of peptides in the time-course experiments were determined in Spectrum Mill using TMT reporter ion intensity ratios from each PSM. TMT reporter ion intensities for the 3 time points were not corrected for isotopic impurities because the intervening 127, 129, and 131 labels were not included. Each peptide-level TMT ratio was calculated as the median of all PSMs contributing to that peptide. PSMs were excluded from the calculation that lacked a TMT label, or had a negative delta forward-reverse identification score (half of all false-positive identifications). Intensity values for each time point were normalized to the 12h time point.

For whole proteome data MS/MS spectra were excluded from searching if they did not have a precursor MH+ in the range of 600-6000, had a precursor charge >5, had a minimum of <5 detected peaks, or failed the spectral quality filter with a sequence tag length >0 (*i.e*., minimum of 2 masses separated by the in-chain masses of 1 amino acid) based on ESI-QEXACTIVE-HCD-v4-30-20 peak detection. Similar spectra with the same precursor m/z acquired in the same chromatographic peak were merged. MS/MS search parameters included: ESI-QEXACTIVE-HCD-v4-30-20 scoring parameters; Trypsin allow P specificity with a maximum of 4 missed cleavages; fixed modification: carbamidomethylation of cysteine and selenocysteine; variable modifications: oxidation of methionine, deamidation of asparagine, acetylation of protein N-termini, pyroglutamic acid at peptide N-terminal glutamine, and pyro-carbamidomethylation at peptide N-terminal cysteine; precursor mass shift range of −18 to 64 Da; precursor mass tolerance of ±20 ppm; product mass tolerance of ± 20 ppm, and a minimum matched peak intensity of 30%. Peptide spectrum matches (PSMs) for individual spectra were automatically designated as confidently assigned using the Spectrum Mill auto-validation module to apply target-decoy based FDR estimation at the PSM level of <1.0% FDR. Protein level data was summarized by top uses shared (SGT) peptide grouping and non-human contaminants were removed. SARS-CoV-2 derived proteins were manually filtered to include identifications with >6% sequence coverage and at least 2 or more unique peptides. Postfiltering, intensity-based absolute quantification (iBAQ) was performed on the whole proteome LC-MS/MS as described in (Schwanhäusser et al., 2011). Briefly, iBAQ values were calculated as follows: log10(totalIntensity/numObservableTrypticPeptides), the total precursor ion intensity for each protein was calculated in Spectrum Mill as the sum of the precursor ion chromatographic peak areas (in MS1 spectra) for each precursor ion with a peptide spectrum match (MS/MS spectrum) to the protein, and the numObservableTrypticPeptides for each protein was calculated using the Spectrum Mill Protein Database utility as the number of tryptic peptides with length 8 - 40 amino acids, with no missed cleavages allowed. Of note, S coverage was 55% in the HEK293T and 44% in the A549 post 24 hour fractionated data, which may be due to the high levels of glycosylation. Lower peptide coverage may lead to underestimation of S protein in our data. Both log10 transformed total intensity and iBAQ values were median normalized by subtracting sample specific medians and adding global medians for each abundance metrics and reported in Supplemental Table S3. For the two-sample T-test analyses shown in Figure 5E,F, only proteins observed across all fractionated uninfected and 24 hr SARS-CoV-2 post infection samples were considered (Fig 5E: n~5,000 protein, Table S3; Figure 5F: n~120 proteins, Table S3).

### Validation of peptide identifications

Peptide identifications were validated using synthetic peptides. Synthetic peptides were obtained from Genscript, at purity >90% purity and dissolved to 10 mM in DMSO. For LC-MS/MS measurements, peptides were pooled and further diluted with 0.1% FA/3% ACN to load 120 fmol/μl on column. One aliquot of synthetic peptides was also TMT labeled as described above. LC-MS/MS measurements were performed as described above. For plots, peak intensities in the experimental and the synthetic spectrum were normalized to the highest peak.

### RNA-Seq

A549 and HEK293T cells were seeded into 6-well plates at a density of 5 x 105 cells per well (one well per condition). After 11-24 hr, the cells were infected with SARS-CoV-2 at MOI of 3. At 12, 18 and 24hr post-infection, the cells were lysed in Trizol (Thermo, 15596026) and the total RNA was isolated using standard phenol chloroform extraction. Standard Illumina TruSeq Stranded mRNA (LT) was performed using 500 ng of total RNA (illumina, FC-122-2101). Oligo-dT beads were used to capture polyA-tailed RNA, followed by fragmentation and priming of the captured RNA (8 minutes at 94°C). Immediately first strand cDNA synthesis was performed. Second strand cDNA synthesis was performed using second strand marking master and DNA polymerase 1 and RNase H. cDNA was adenylated at the 3’ ends followed immediately by RNA end ligation single-index adapters (AR001-AR012). Library amplification was performed for 12-15 cycles under standard illumina library PCR conditions. Library quantitation was performed using Agilent 2200 TapeStation D1000 ScreenTape (Agilent, 5067-5582). RNA sequencing was performed on the NextSeq 550 System using a NextSeq V2.5 High Output 75 cycle kit (illumina, 20024906) or 150 cycles kit (illumina, 20024907) for paired-end sequencing (70nt of each end).

### RNA sequencing reads alignment

Sequencing reads were mapped to SARS-CoV-2 genome (RefSeq NC_045512.2) and human transcriptome (Gencode v32). Alignment was performed using Bowtie version 1.2.2 (Langmead et al., 2009) with a maximum of two mismatches per read. The fraction of human and viral reads was determined based on the total number of reads aligned to either SARS-CoV-2 or human transcripts. The raw RNA sequencing data generated in this study have been submitted to the Gene Expression Omnibus (GEO; https://www.ncbi.nlm.nih.gov/geo/) under accession number GSE159191.

### HLA-I antigen presentation prediction

HLAthena, a prediction tool trained on endogenous LC-MS/MS-identified epitope data, was used to predict HLA class I presentation for all unique 8-11mer SARS-Cov-2 peptides across 31 HLA-A, 40 HLA-B and 21 HLA-C alleles(Sarkizova et al., 2020).

### HLA allele frequencies and coverage estimates

World frequencies of HLA-A, -B, and -C allele in Table S6B are based on a meta-analysis of high-resolution HLA allele frequency data describing 497 population samples representing approximately 66,800 individuals from throughout the world(Solberg et al., 2008), downloaded from http://pypop.org/popdata/2008/byfreq-A.php.html, http://pypop.org/popdata/2008/byfreq-B.php.html, http://www.pypop.org/popdata/2008/byfreq-C.php. Subpopulation frequencies for AFA, API, EUR, HIS, and USA were obtained from supplementary data in (Poran et al., 2020a).

The cumulative phenotypic frequency (CPF) of peptides was calculated using *CPF* = 1 – (1 – ∑*_i ∈ C_ p_i_*)^2^, assuming Hardy-Weinberg proportions for the HLA genotypes(Dawson et al., 2001), where *p_i_* is the population frequency of the *i*^th^ alleles within a subset of HLA-A, -B, or C alleles, denoted *C*. Coverage across HLA-A, -B, and -C alleles was calculated similarly: *CPF* = 1 – (1 – ∑*_i ∈ A_ p_i_*)^2^ * (1 – ∑*_i ∈ B_ p_i_*)^2^ * (1 – ∑*_i ∈ C_ p_i_*)^2^, where A, B, and C denote a subset of HLA-A, -B, and/or -C alleles for which the coverage is computed, as recently done in (Poran et al., 2020a).

