## Supplemental Data for "SARS-CoV-2 infected cells present HLA-I peptides from canonical and out-of-frame ORFs"

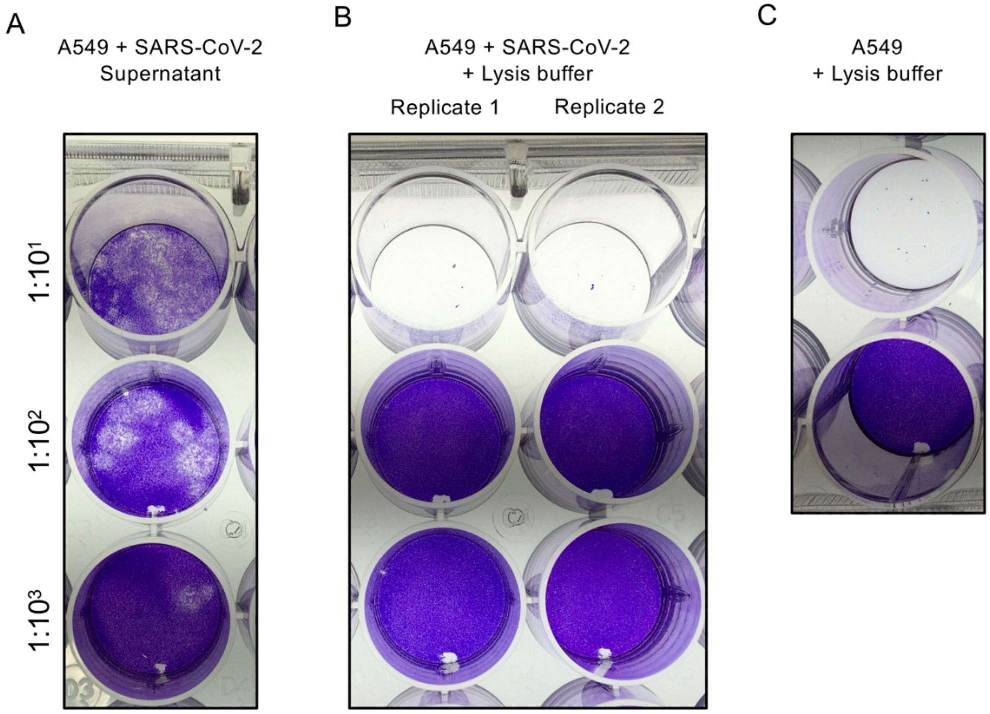


**Figure S1. SARS-CoV-2 inactivation for HLA-IP experiments**

A549 cells were infected with SARS-CoV-2 at MOI of 3 for 24 hours and virus inactivation was determined by plaque assay. 10-fold serial dilutions were prepared in Opti-MEM and used to infect Vero cells in a 24-wells plate. **(A)** Cultured media of infected A549 cells. As expected, plaques were observed in all three dilutions, confirming the presence of infectious virus. **(B)** Infected A549 cells were treated with a lysis buffer containing 1.5% Triton-X and Benzonase. Lysates were incubated for 3 hours in the BSL3 prior dilution for plaque assay. When adding the 1:10 dilution, cells died immediately due to the relatively high Triton-X concentration. No plaques were observed in the $1{:}10^2$ and $1{:}10^3$ dilutions. **(C)** Same as (B) but for non-infected A549 cells. Here too, 1:10 dilution leads to immediate cell death confirming that the observed cell death in the infected sample was not due to the virus. As expected, no plaques were observed.

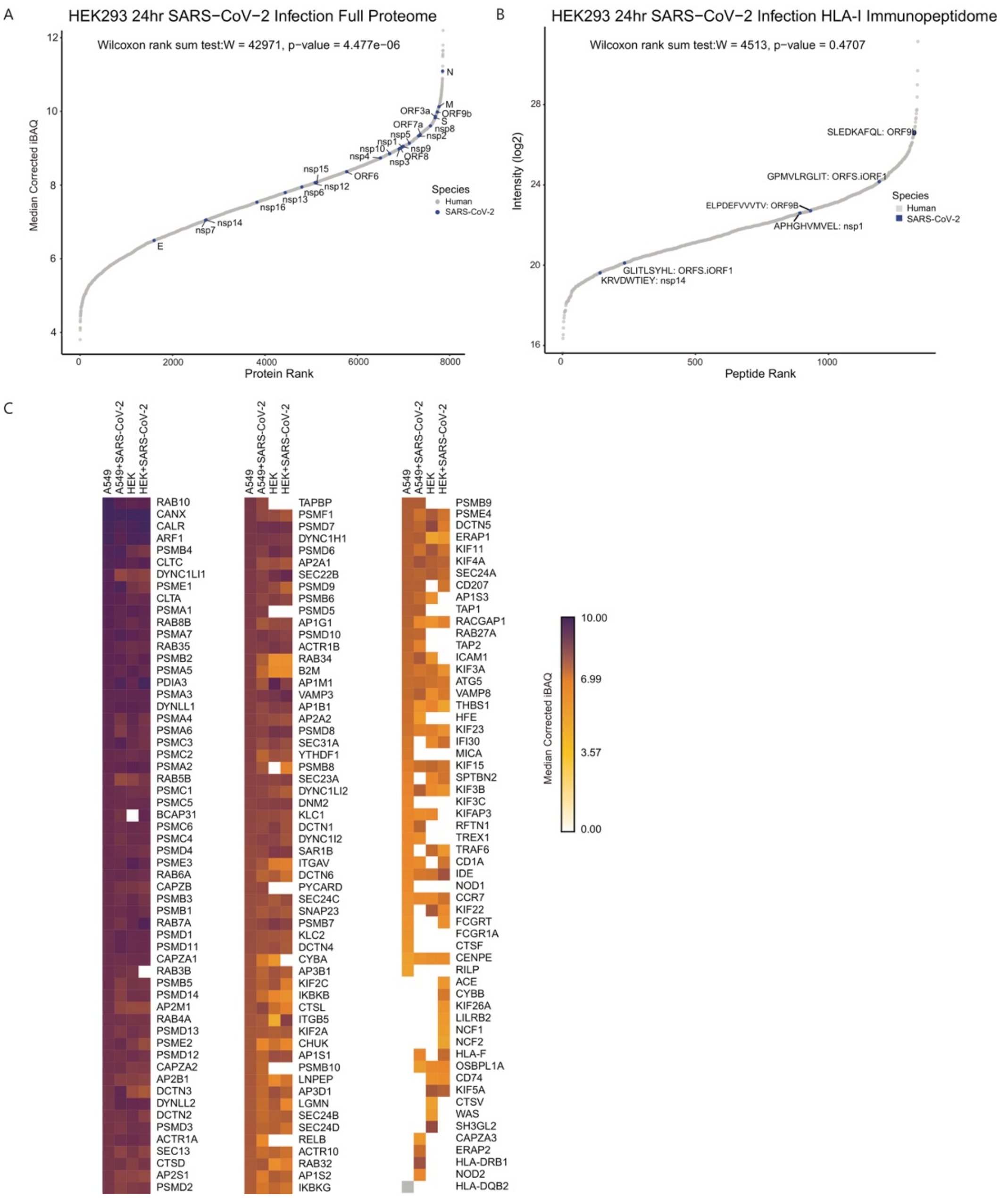


**Figure S2. SARS-CoV-2 peptides abundance and antigen presentation pathway proteins in infected cells**

**(A)** Rank plot of the iBAQ value for each human (gray) and SARS-CoV-2 (blue) protein detected in HEK293T cells 24 hours post infection. SARS-CoV-2 proteins are annotated with their respective gene names. **(B)** Similar rank plot of (A) but for observed HLA-I peptides and their intensities. Peptides mapped to SARS-CoV-2 are annotated with their respective amino acid sequence and source protein gene name. **(C)** Heatmap of iBAQ values for antigen presentation pathway proteins observed across uninfected and 24 hours post infection in A549 and HEK293T cells.

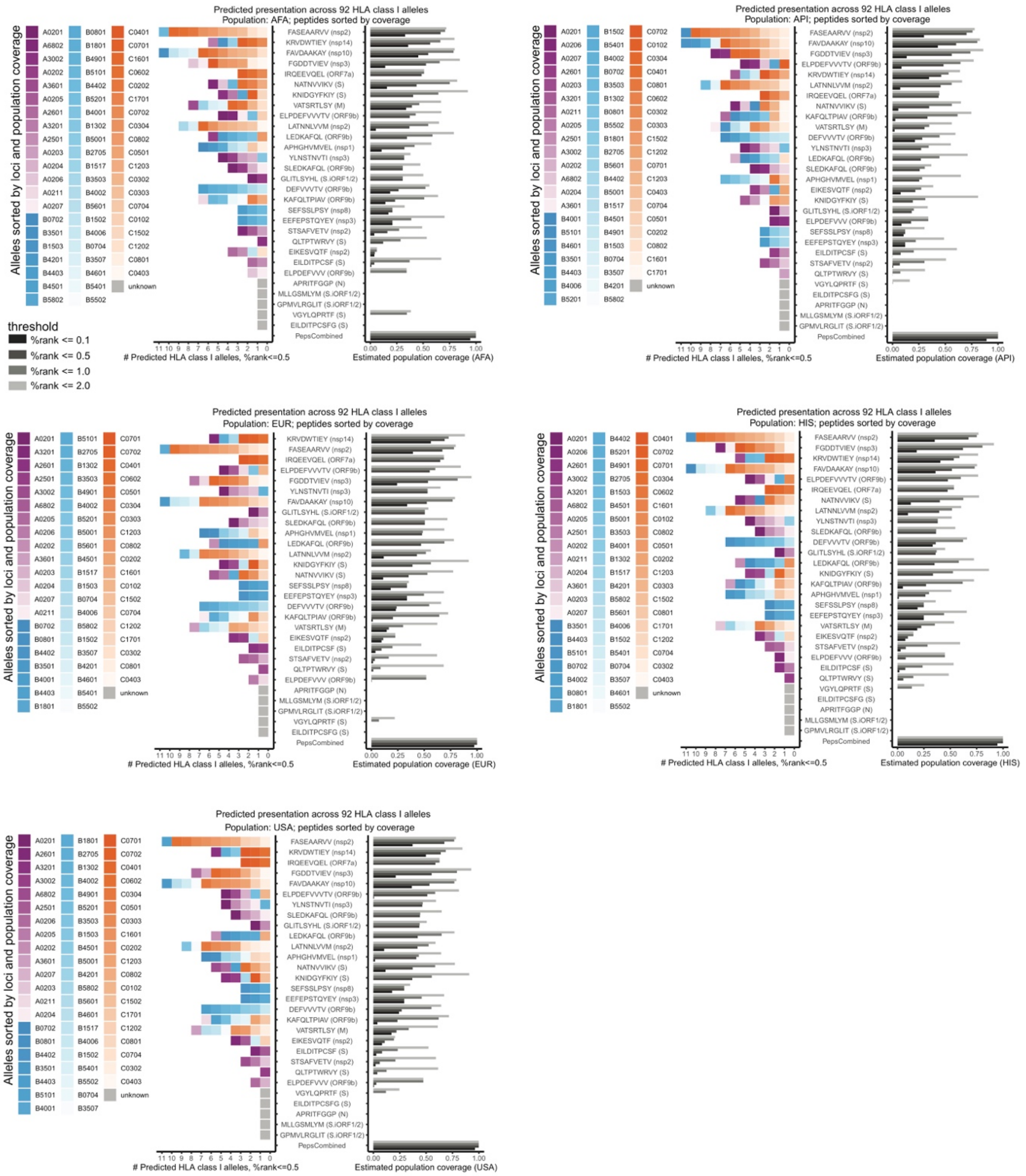


**Figure S3. Population coverage estimates of MS-identified SARS-CoV-2 HLA-I peptides.** Similar to Fig 6, HLAthena predictions for 92 HLA-I alleles using percentile rank cutoff values of 0.1, 0.5, 1, and 2% were used to show the number of alleles and estimated coverage for each LC-MS/MS-observed SARS-CoV-2 peptides across **(A)** AFA, **(B)** API, **(C)** EUR, **(D)** HIS, and **(E)** USA populations. Alleles are colored and ordered according to loci and the corresponding population frequency (high to low color intensity). Peptides are ordered according to their estimated coverage at %rank cutoff of 0.5.

### SUPPLEMENTAL TABLE LEGENDS

**Table S1. HLA-I peptidomics.** Peptide identifications from HLA-I immunoprecipitation experiments in A549 and HEK293T cells +/- SARS-CoV-2 after filtering for contaminants and length 8-11. For TMT time course experiments, peptides were filtered for quantitative values and TMT ratios to the 12h time point were median normalized.

**Table S2. SARS-CoV-2 protein abundance and presentability.** (A) SARS-CoV-2 protein expression values as determined by whole proteome measurements in A549 and HEK293T cells and Ribo-Seq translation measurements that were previously published for Vero cells (Finkel et al., 2020b). iBAQ - intensity-based absolute quantification. (B) HLA-I presentability estimates of SARS-CoV-2 ORFs based on HLAthena predictions.

**Table S3. Whole proteome data.** Data associated with whole proteome analyses of A549 and HEK293T cells +/- SARS-CoV-2 at 12hr, 18hr, and 24 hr time points post infection. Two sample t-test results (Figure 5E, F) are summarized for all proteins observed across both cell types and proteins involved in ubiquitination pathways.

**Table S4. HLA-I predictions and Coverage.** (A) HLA-I presentation prediction of SARS-CoV-2 epitopes per cell line. (B) HLA-I alleles frequencies. (C) Estimated population coverage of MS-detected SARS-CoV-2 epitopes based on predictions across 92 HLA-I alleles.
